# Label-free long-term measurements of adipocyte differentiation from patient-driven fibroblasts and quantitative analyses of in situ lipid droplet generation

**DOI:** 10.1101/2024.01.06.574492

**Authors:** Eun Young Jeong, Hye-Jin Kim, Sumin Lee, Yongkeun Park, Young Mo Kim

**Affiliations:** Department of Orthopedic Surgery, College of Medicine, Chungnam National University, Daejeon 34134, Republic of Korea; Tomocube Inc., Daejeon 34109, Republic of Korea; Department of Physics, Korea Advanced Institute of Science and Technology (KAIST), Daejeon 34141, Republic of Korea; KAIST Institute for Health Science and Technology, KAIST, Daejeon 34141, Republic of Korea

## Abstract

The visualization and tracking of adipocytes and their lipid droplets (LDs) during differentiation are pivotal in developmental biology and regenerative medicine studies. Traditional staining or labeling methods, however, pose significant challenges due to their labor-intensive sample preparation, potential disruption of intrinsic cellular physiology, and limited observation timeframe. This study introduces a novel method for long-term visualization and quantification of biophysical parameters of LDs in unlabeled adipocytes, utilizing the refractive index (RI) distributions of LDs and cells. We employ low-coherence holotomography (HT) to systematically investigate and quantitatively analyze the 42-day redifferentiation process of fat cells into adipocytes. This technique yields three-dimensional, high-resolution refractive tomograms of adipocytes, enabling precise segmentation of LDs based on their elevated RI values. Subsequent automated analysis quantifies the mean concentration, volume, projected area, and dry mass of individual LDs, revealing a gradual increase corresponding with adipocyte maturation. Our findings demonstrate that HT is a potent tool for non-invasively monitoring live adipocyte differentiation and analyzing LD accumulation. This study, therefore, offers valuable insights into adipogenesis and lipid research, establishing HT and image-based analysis as a promising approach in these fields.

## 1. Introduction

Knee osteoarthritis (KOA) is a progressive, degenerative joint disorder marked by cartilage degradation and concurrent bone remodeling, primarily due to mechanical overloading. Recent insights into KOA pathogenesis highlight the significant roles of proinflammatory and immune-related signaling pathways alongside biomechanical factors [1]. The infrapatellar fat pad (IFP), or Hoffa’s fat pad, an adipose tissue structure within the knee’s anterior compartment [2], is increasingly recognized for its biomechanical and pathophysiological roles in KOA [3-9]. IFP is implicated in immune cell infiltration and the production of proinflammatory cytokines (e.g., IFNγ, TNFα, IL1β) and cartilage-degrading molecules (e.g., MMPs) [10-12].

Adipocytes, or fat cells, are known to contribute significantly to osteoarthritis progression through their secretion of adipokines and cytokines, which regulate inflammatory responses [13-15]. Adipose tissue comprises various cell types, including preadipocytes, mature adipocytes, endothelial, and immune cells [16]. Notably, the differentiation process of adipocytes within the IFP is critical in KOA progression. The IFP undergoes morphological and cellular changes during KOA, including the proliferation and differentiation of adipocytes into fibroblast-like dedifferentiated fat (DFAT) cells. These DFAT cells are implicated in producing pro-inflammatory mediators and matrix metalloproteinases that contribute to cartilage degradation [17]. Previous studies suggest that both the rate and extent of adipocyte differentiation are altered in KOA [18]. The inflammatory environment within the KOA joint can accelerate adipocyte differentiation, leading to increased adipokine production, which further exacerbates inflammation and cartilage degradation [19]. Understanding the dynamics of this differentiation process could provide key insights into disease progression and identify potential therapeutic targets.

Morphologically, mature adipocytes, preadipocytes, and DFAT cells display distinct differences, particularly in the number and size of lipid droplets (LDs). The larger and less numerous LDs in mature adipocytes underscore their lipid storage specialization, contrasted by the numerous smaller droplets in preadipocytes and variable droplet characteristics in DFAT cells [20]. This differentiation process, particularly the behavior of lipid droplets within these cells, could inform strategies for diagnosing KOA and designing targeted treatments that modulate adipogenic activity within the IFP, influencing overall disease pathology and patient outcomes [21].

LDs, enclosed by a monolayer of phospholipids and proteins, are the primary lipid-storing organelles in cells, especially adipocytes [22-24]. Adipocyte differentiation, or adipogenesis, is marked by increasing LD accumulation, a process central to lipogenesis [25]. LD dynamics are also linked to metabolic conditions, including cancer, obesity, and diabetes, making their analysis crucial for understanding metabolic health [26]. Recently, various methods have been developed to accurately monitor fat cell function through morphological analysis of LD parameters [27-29]. Traditional methods for tracking and quantifying LDs, such as Nile red, Oil-red O, BODIPY, and other fluorescent dyes [17, 20, 30-34], provide only static snapshots and may disrupt live cell monitoring due to the requirement for cell fixation [35]. Conversely, phase-contrast and non-invasive imaging tools enable dynamic LD tracking but lack the ability to observe live cells over extended periods [36].

To investigate the dynamics of LDs in live cells while preserving their physiological conditions, several label-free imaging techniques have been utilized [37]. These include coherent anti-Stokes Raman spectroscopy (CARS) [38], stimulated Raman scattering (SRS) [39], and quantitative phase imaging (QPI) [40]. technique to monitor lipid distribution and dynamics in live cells [41] Additionally, autofluorescence lifetime imaging (FLIM) has emerged as a label-free imaging method. FLIM provides critical insights into lipid dynamics and cellular metabolic states, which are particularly relevant in understanding adipogenesis in osteoarthritis [41].

QPI represents a significant advancement in cellular imaging by utilizing variations in the optical path length of light transmitted through transparent samples to generate image contrast. This contrast is particularly pronounced in cellular components such as LDs, which have distinct refractive indices (RIs) due to their composition. Among the various forms of QPI, holotomography (HT) has emerged as a specialized technique that capitalizes on these RI differences to detect and analyze LDs within cell [42-46]. Unlike traditional imaging methods, HT provides a non-invasive approach to visualize and quantify LDs in three dimensions, offering critical data on their volume, density, and dynamic changes over time without the need for dyes or labels that could alter cellular physiology. This capability makes HT particularly valuable for studies in adipocyte biology, where understanding the intricate dynamics of LDs is essential for elucidating processes like lipid metabolism and adipogenesis. The introduction of HT in LD imaging not only enhances our ability to observe these cellular structures in their native state but also opens new avenues for exploring metabolic disorders and potential therapeutic targets in adipocyte-related diseases. By integrating this technology into our research, we aim to provide deeper insights into LD behavior and its implications for disease progression and treatment.

In this study, we demonstrate long-term measurements of adipocyte differentiation from patient-driven fibroblasts utilizing low-coherence HT. Employing quantitative imaging analysis algorithms, we non-invasively detect and analyze lipid droplets within intact adipocytes over forty-two days of differentiation processes. This approach facilitates a detailed examination of the morphological attributes of mature adipocytes as they redifferentiate from DFAT cells obtained from IFP of patients undergoing osteoarthritis surgery. Our analysis yields comprehensive evaluations of droplet volumes, lipid concentrations, and contents throughout a 42-day observation period. By integrating these insights into the broader narrative of KOA progression, we aim to provide an understanding of adipocyte differentiation’s role in KOA pathology and treatment. The methodology presented herein promises to be a powerful tool for elucidating the processes of adipogenesis and LD accumulation across diverse cell types.

## 2. MATERIALS AND METHODS

### Cell isolation and culture

Isolation and expansion of human primary preadipocytes were carried out using tissues from IFP obtained from patients with osteoarthritis (OA), with slight modifications to previously described methods [6]. In brief, we processed approximately 1 g of adipose tissue by mincing and enzymatic dissociation in 0.1% (w/v) collagenase type II (Worthington Biochemical Corporation, Lakewood, NJ) at 37°C for 2 hours with gentle shaking. The resulting cell suspension was filtered through a 1000 μm nylon mesh to separate cells from unwanted stromal fragments and tissue residues. The mature adipocytes, which floated to the surface, were collected and centrifuged at 135 g for 3 minutes. These isolated adipocytes were then cultured in 25 cm^2^ flasks (Corning^®^) filled with DMEM (GIBCO^®^) supplemented with 10% (v/v) fetal calf serum (FCS, Hyclone) and 1% (v/v) antibiotics, and maintained at 37°C in a 5% CO2 atmosphere.

For the ceiling culture technique, the T25 flasks were oriented with the adhesive surface facing upwards to allow the buoyant, lipid droplet-rich adipocytes to attach to the flask’s inner ceiling. After 1 to 2 weeks, the medium was discarded, the flasks inverted, and fresh medium added just enough to cover the flask’s bottom. Upon reaching confluency, the cells were passaged and utilized for subsequent experiments. This study is approved by the Chungnam University Research Review Board (IRB file No. 2020-05-009-012).

### Differentiation assays

To induce adipogenic differentiation, human preadipocytes were seeded in 12-well plates (Cellvis) and cultured to confluency. Adipogenic induction was then initiated using AdipoLife^®^ Initiation Medium (LifeLine Cell Tech) for 6 days, with the medium refreshed bi-daily. From Day 6 to Day 45, the cells were maintained in AdipoLife^®^ Maintenance Medium (LifeLine Cell Tech) to facilitate further differentiation. At specified time points during the differentiation process, total RNA was harvested for subsequent quantitative real-time RT-PCR analysis.

### Holotomography

Three-dimensional RI tomograms of individual cells were captured using low-coherence HT system. HT, also known as a 3D QPI technique, reconstructs 3D RI tomogram of an unlabeled sample from multiple 2D optical images with various illumination modulations [40, 47]. Due to its label-free tomographic imaging and high contrast for lipid droplets, HT has been utilized for detecting LDs in unlabeled cells [42, 48], biomolecular condensation [49, 50], and biophysics [51, 52].

The used HT system (HT-X1, Tomocube Inc., Daejeon, Republic of Korea) utilizes an LED with a central wavelength of 450 nm for illumination, providing stable, label-free cellular imaging with minimal speckle noise. Wavefront modulation is achieved through a digital micromirror device (DMD) situated at the pupil plane, paired with a specially designed condenser lens (NA 0.72) that features a long working distance of 30 mm. Coupled with a motorized module and a high numerical aperture objective lens (NA 0.95, UPLXAPO40X, Olympus), the system captures multiple intensity stacks from thick specimens under variously modulated illuminations. These stacks are processed to reconstruct the sample’s 3D RI tomogram. Further methodological details, including intensity-based field retrieval, tomogram reconstruction via deconvolution, and regularization algorithms for the missing cone problem, are elaborated in other publications [53-56]. The image acquisition time for a field-of-view (FOV) of 150 μm × 150 μm × 50 μm field of view is approximately three seconds, with a reconstruction time of 50 seconds. Theoretical resolution reaches 160 nm laterally and 1 μm axially [40].

### Time-course holotomography imaging

Prior to the introduction of adipogenesis initiation medium (AIM), DFAT in their culture vessel were mounted on the HT-X1 system. The integrated stage top incubation chamber (TOKAI HIT, Japan) within the holotomography setup ensures precise temperature control at 37°C and a 5% CO_2_ atmosphere, optimal for extended live-cell imaging. Ten distinct locations within two wells were systematically selected for automated imaging using TomoStudioX software (Tomocube Inc., Daejeon, Republic of Korea). Each site was imaged with a FOV of 400 μm × 400 μm and a z-depth of 60 μm. Following initial HT imaging on day 1, the culture vessel was temporarily removed from the system, and half of the medium was refreshed with AIM.

The DFAT cells were then cultured in a 5% CO_2_ incubator for a 7-day maturation period. Subsequently, the vessel was repositioned onto the HT system, and the previously recorded ten positions were revisited to capture images under identical conditions. Post Day 7 imaging, the vessel was removed, and half of the culture medium was replaced with adipogenesis maintenance medium (AMM). This cycle of imaging at intervals of 3-4 days followed by medium renewal was continued for five weeks to facilitate and monitor the differentiation process.

### Quantitative image analysis of Lipid Droplets in Adipocytes

From the reconstructed RI tomogram of unlabeled adipocytes, various biophysical parameters of individual LDs were retrieved. LDs were segmented using the Adipocytes LD Segmentation pipeline in the TomoAnalysis software (Tomocube Inc., Daejeon, Republic of Korea). The segmentation process was carried out in two steps. Firstly, the images were thresholded using a RI value of 1.37 and subsequently post-processed with an opening algorithm to identify large LDs. Secondly, to detect small LDs that were still in the growth phase, the images were filtered using the Tophat method and then subjected to a threshold. The combination of these two steps effectively segmented LDs from the HT images

The volumes of individual LDs were calculated by counting the number of voxels corresponding to LDs, segmented from their high RI values. Compared to other subcellular organelles, LDs exhibit distinctly higher RI values [42]. The mean RI values of the individual LDs were calculated by averaging RI values corresponding to LDs. The dry mass of lipids in individual LDs were calculated by using a linear relationship between the RI and the concentration of biomolecules [57, 58]. The used proportional value, the RI increment for LDs, was 0.135 mL/g for LDs [42]. The projected areas for individual LDs were calculated by projecting the volumes of LDs along the axial direction, from which the maximal areas of LDs were retrieved.

### Immunofluorescence imaging

For immunofluorescence staining, cells were fixed with 4% paraformaldehyde (Sigma– Aldrich, US) for 15 minutes, rinsed with PBS, and then stained for 10 minutes using LipidSpot™ Lipid Droplet Stain (Cambrex, US) to highlight lipid droplets. After a subsequent PBS wash, cells underwent DNA staining with 1 mg/ml DAPI (Sigma–Aldrich). Multichannel 3D fluorescence imaging was performed in conjunction with using the HT system (HT-X1, Tomocube Inc., Republic of Korea).

### RNA extraction and Real-time quantitative RT-PCR

Total RNA was extracted from cells utilizing an RNA extraction Kit, according to the manufacturer’s instructions. The isolated RNA was then reverse-transcribed into cDNA using a cDNA Reverse Transcription Kit (TOYOBO, Japan). The synthesized cDNA served as a template for RT-PCR amplification, employing PCR Master Mix (Bionner, Daejeon, Republic of Korea) and gene-specific primers. PCR product band intensities were quantified using Image-Pro Plus 6.0 software, and expression levels were normalized to the internal control gene, GAPDH. Quantitative real-time RT-PCR analyses were conducted with the SYBR Green RT2 kit (Bionner, Daejeon, Republic of Korea), using a CFX96 C1000 thermal cycler (Bio-Rad, US). GAPDH was also quantified as a housekeeping gene in each sample to standardize the results. The relative expression of target genes was calculated using the ΔΔCt method and presented as fold change normalized to GAPDH expression.

### Statistical analysis

Data are presented as the mean ± standard deviation (SD) for three independent experiments (N = 3). Statistical significance was evaluated using a two-tailed t-test for multiple comparisons, with significance thresholds set at ^***^ *p* < 0.001 and ^**^ *p* < 0.01. All analyses were performed using Prism 9 (GraphPad Software Inc.), which facilitated the identification of significant quantitative differences between comparative groups.

## 3. RESULTS

### Label-free observation of whole redifferentiation process of DFAT cells

Utilizing a label-free HT imaging approach, we tracked the entire redifferentiation process of DFAT cells derived from human IFP [59]. The 3D imaging capabilities of our HT instrument enabled us to observe dynamic morphological changes in adipocytes over the 42-day differentiation period. These mature adipocytes, isolated from OA patients’ knee joints post-surgery, underwent a ceiling culture method to facilitate dedifferentiation (Figure 1A). The ability to capture and analyze these changes in three dimensions was crucial for understanding adipocyte morphology, particularly in terms of lipid droplet growth and distribution within the cells. Within one week of this unique culture, we observed a marked morphological shift from the characteristic rounded adipocyte form to a more fibroblast-like cell structure. This period was characterized by the cells adopting a flattened morphology, developing membranous extensions, and displaying increased motility. Upon reaching a high level of confluency, these cells were then passaged for use in subsequent experiments. These fibroblast-like cells, now referred to as DFAT cells, represent a transitional state in the redifferentiation process (Figure 1B)

**Figure 1:**
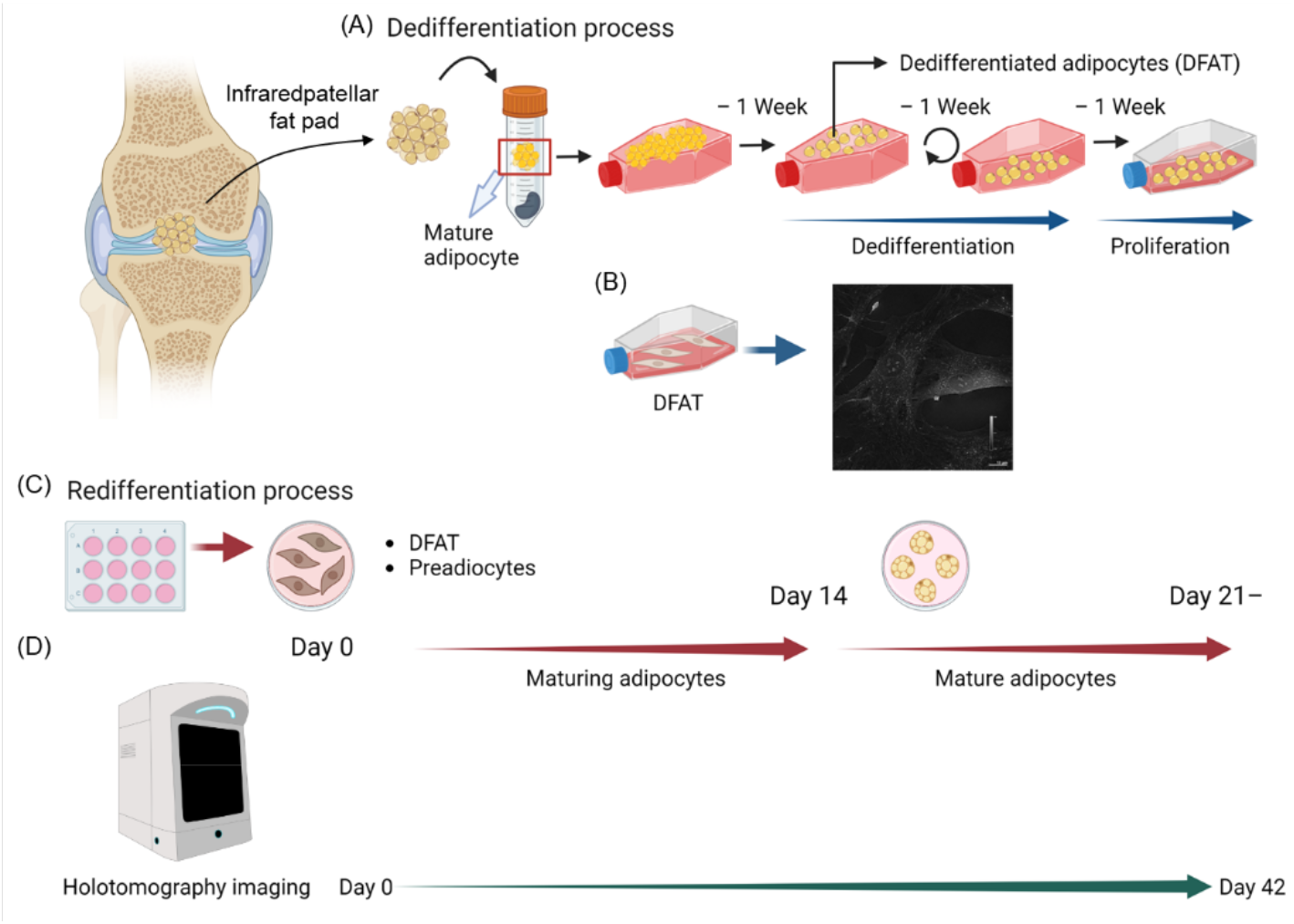
Schematic Representation of Adipocyte Dedifferentiation and Redifferentiation Tracked by Holotomography Imaging. (A) Dedifferentiation process: Harvested mature adipocytes from the infrapatellar fat pad undergo a ceiling culture method for dedifferentiation into dedifferentiated adipocytes (DFAT) over three weeks, with each week marked by changes in cell morphology and proliferation. (B) Holotomographic image of DFAT cells highlighting the characteristic morphological features, with scale bar indicating size. (C) Redifferentiation process: Starting from day 0 with DFAT and preadipocytes, the transition is depicted through day 14, showing maturing adipocytes, and continues until mature adipocytes are formed from day 21 onwards. (D) Holotomography imaging timeline: Depiction of the non-invasive imaging schedule using holotomography from day 0 to day 42 to monitor the morphological changes during the redifferentiation process.

Over the course of 1-2 weeks in culture, the attached fat cells progressively lost their unilocular lipid droplet structure, indicative of dedifferentiation. The 3D imaging feature was instrumental in performing quantitative volume measurements of lipid droplets, which were critical for assessing lipid accumulation and understanding adipogenesis. This was visually corroborated by the diminishing lipid content, as the cells transitioned from a state of lipid storage to a more proliferative and less differentiated phenotype (Figure 1C). The HT system allowed for precise spatial distribution and compartmentalization analysis of lipid droplets within adipocytes, providing insights into their interaction with other cellular structures, which is important for elucidating their role in cellular metabolism and inflammatory responses. We continuously imaged the same areas every three days, capturing the nuanced stages of redifferentiation as the DFAT cells matured back into adipocytes (Figure 1D).

The redifferentiation process was successfully visualized using the HT, circumventing the need for traditional fluorescence labeling methods. The HT images provided a high-resolution, three-dimensional view of the adipocytes, highlighting the morphological features characteristic of DFAT cells (Figure 2A). These 3D reconstructions provided higher resolution and more accurate representations of cell structures, enhancing the integrity of our data. Overall morphology and RI values of the LDs in DFAT cells are comparable to the previously reported LDs in hepatocyte and form cells [42, 43]. The fluorescence staining further confirmed these observations, with lipid droplets and nuclei distinctly illuminated. The overlay of segmentation masks on fluorescence images validated the accuracy of our label-free detection technique, underscoring the reliability of RI thresholding as a method for lipid droplet quantification (Figure 2B).

**Figure 2:**
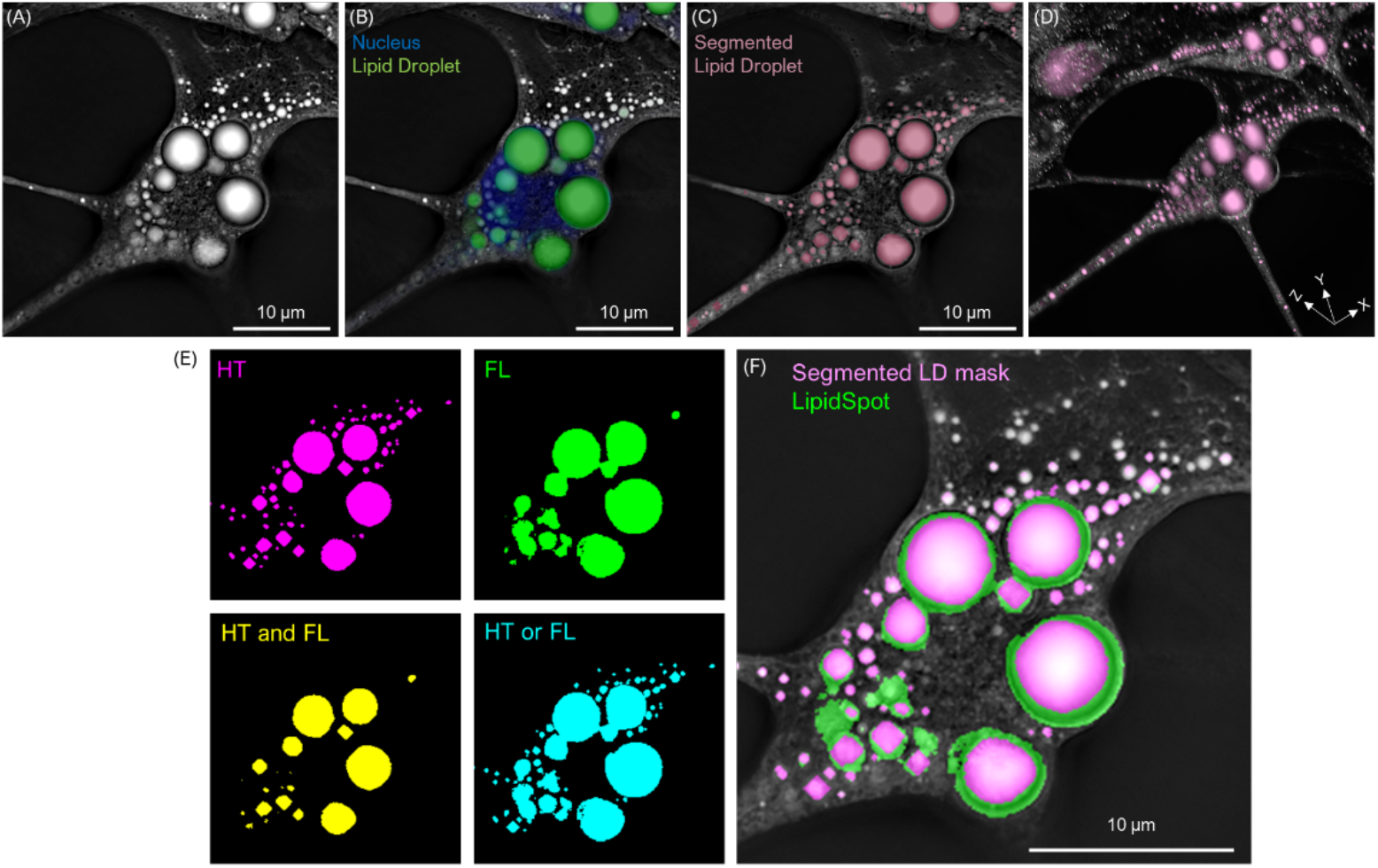
Correlation of Refractive Index with Lipid Droplets in Adipocytes. (A) Holotomography of mature adipocytes with corresponding refractive index analysis. Regions with a refractive index (RI) above 1.37, typically associated with lipid droplets, are evident. (B) Visualization of lipid droplets in mature adipocytes using a specific fluorescent dye (green) for lipid droplet staining and Hoechst (blue) for nucleus, merged with holotomography (grayscale).(C) Comparison and overlay of segmentation masks (identifying regions with a refractive index above 1.37) with the fluorescence-stained lipid droplets, demonstrating precise correspondence.(D) Three-dimensional rendering of the refractive index distribution within adipocytes, highlighting the segmented lipid droplets (pink) in 3D space. (E) Comparisons of the masks generated from HT (top left), fluorescence labeling (FL, top right), the union of HT and FL (bottom left), and the intersection of HT and FL (bottom right). (F) Overlay of HT, the HT-based mask, and the FL-based mask.

To demonstrate systematic analysis capability, the individual LDs were identified with their high RI values (>1.37), a threshold indicative of lipid content (Figure 2C). When these label-free RI assessments were compared to lipid droplets stained with LipidSpot^®^ 488, there was a precise match, underscoring the reliability of RI thresholding as a method for lipid droplet quantification (Figure 2C and 2D). We have performed a quantitative assessment using the Dice similarity coefficient to measure the overlap between labeled and masked lipid droplets in 3D space. This analysis offers a more rigorous validation of our segmentation approach, ensuring a more accurate comparison between the labeled and detected droplets (Figure 2E and 2F). The 3D Dice score and the Structural Similarity Index Measure (SSIM) were calculated as 0.5014 and 0.7655, respectively. The result shows a successful application of HT to monitor and quantify the redifferentiation of adipocytes. The present approach offers an effective way to observe cellular morphology and lipid droplet dynamics in real-time, providing invaluable insights into adipocyte biology and the complex process of redifferentiation.

### Morphological and functional progression of adipocytes in redifferentiation

To further study the redifferentiation of DFAT cells into mature adipocytes over a 42-day period, the physical expansion of lipid droplets was investigated as a key morphological indicator of cell maturation. The 3D imaging capability provided detailed visual evidence of the lipid droplets’ progressive growth within a consistent FOV (Figure 3A). From days 11 through 42, the yellow dashed outlines in Figure 3A illustrated the lipid droplets’ enlargement, correlating with the cells’ functional evolution during the redifferentiation process.

**Figure 3:**
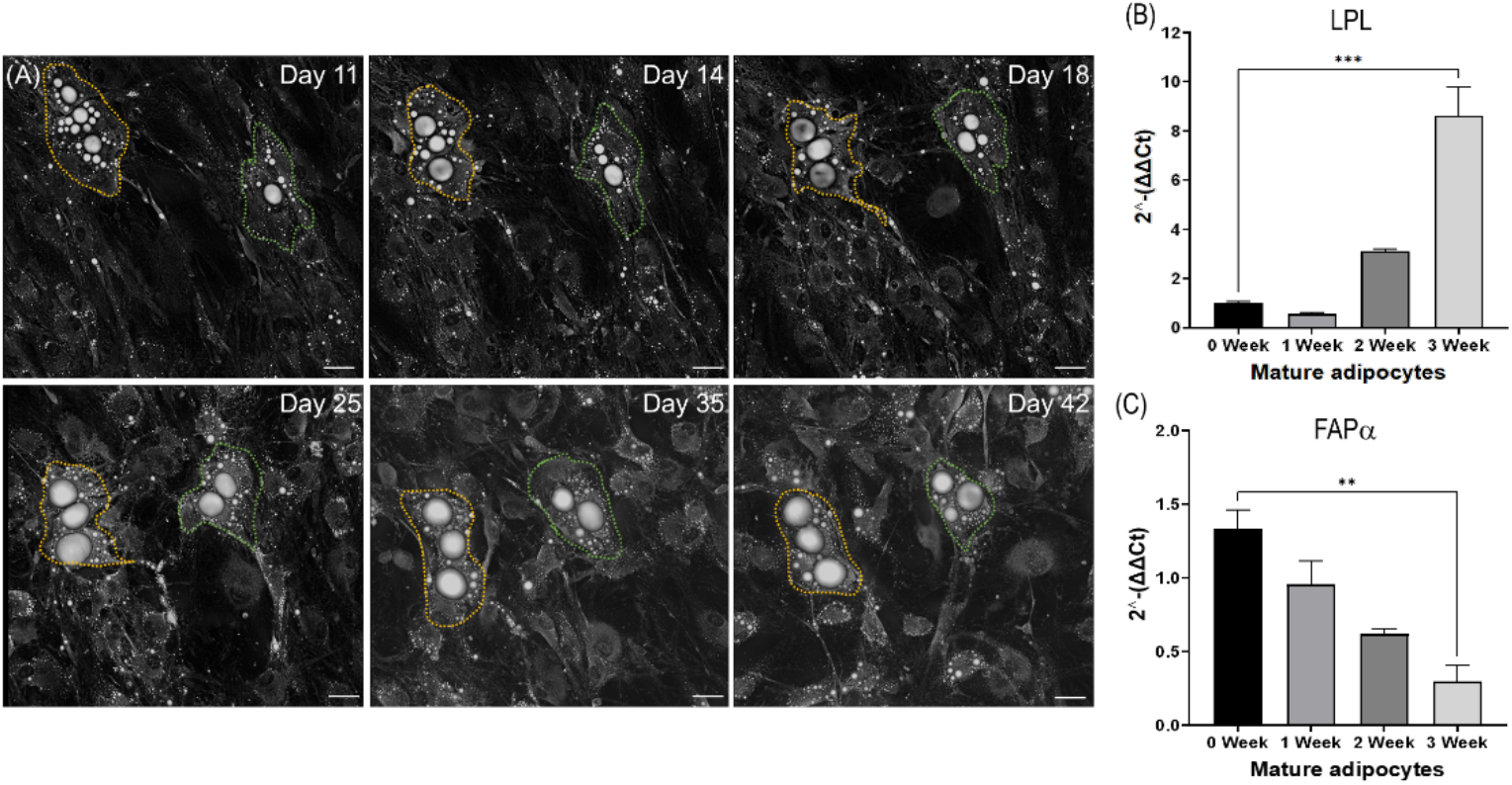
Dynamic Visualization of Adipocyte Redifferentiation and Associated Gene Expression Over a 42-Day Period. (A) Time-lapse holotomography of DFAT cells transitioning to mature adipocytes, captured in the same field. Images from select time points (Days 11, 14, 18, 25, 35, and 42) showcase the progressive enlargement of lipid droplets, delineated by colored dashed outlines, indicative of adipocyte maturation. (B) Quantitative RT-PCR analysis of Lipoprotein Lipase (LPL) and Fibroblast Activation Protein alpha (FAPα) gene expression, a marker for mature adipocyte function, over the course of redifferentiation. The expression levels, presented as fold change (2^-ΔΔCt) relative to day 0, display a significant increase, denoting functional maturation of adipocytes. Data are expressed as mean ± SD (N = 3), with *** indicating P < 0.001, and ** denoting P < 0.01, as determined by a two-tailed t-test. Scale bars represent 40 μm.

This structural maturation was paralleled by significant changes in gene expression related to adipocyte functionality. The RT-PCR analysis revealed a marked upregulation of Lipoprotein Lipase (LPL) as an essential markers of mature adipocyte function and down regulation of Fibroblast Activation Protein alpha (FAPα) which is the marker of fibroblast cell type (Figures 3B and 3C). The expression levels showed the cells obtained characters of adipocytes instead of fibroblast over the study period, with statistical analyses confirming the changes’ significance. These findings underscore the functional maturation of adipocytes, indicating a return to a state capable of active lipid metabolism and storage.

However, during the first week of the study, LPL expression displayed a noticeable decrease before its subsequent increase. This decrease likely corresponds to the early phase of adipocyte differentiation, where early differentiation markers can fluctuate as the cells undergo significant morphological and functional changes. This phase may involve transient downregulation of certain genes, including LPL, before the cells reach a more stable state of differentiation. This pattern is consistent with findings in other studies of cellular differentiation, where similar dynamics have been observed.

### Quantitative Assessment of Lipid Droplet Evolution

In conjunction with morphological observations, a quantitative assessment of lipid droplet parameters provided insight into the biophysical changes accompanying adipocyte redifferentiation (Figure 4). Ten randomly selected FOV were imaged tri-weekly, and parameters of volume, mean RI, dry mass, and projected areas of lipid droplets were measured. In our methodology, LDs within adipocytes were segmented based on their high RI values (see Methods). This segmentation involved a two-step process for comprehensive identification of LDs at various stages of development. Initially, the HT images were subjected to a thresholding process based on a RI value of 1.37, followed by the application of an opening algorithm. This step was primarily aimed at isolating larger, mature LDs.

**Figure 4:**
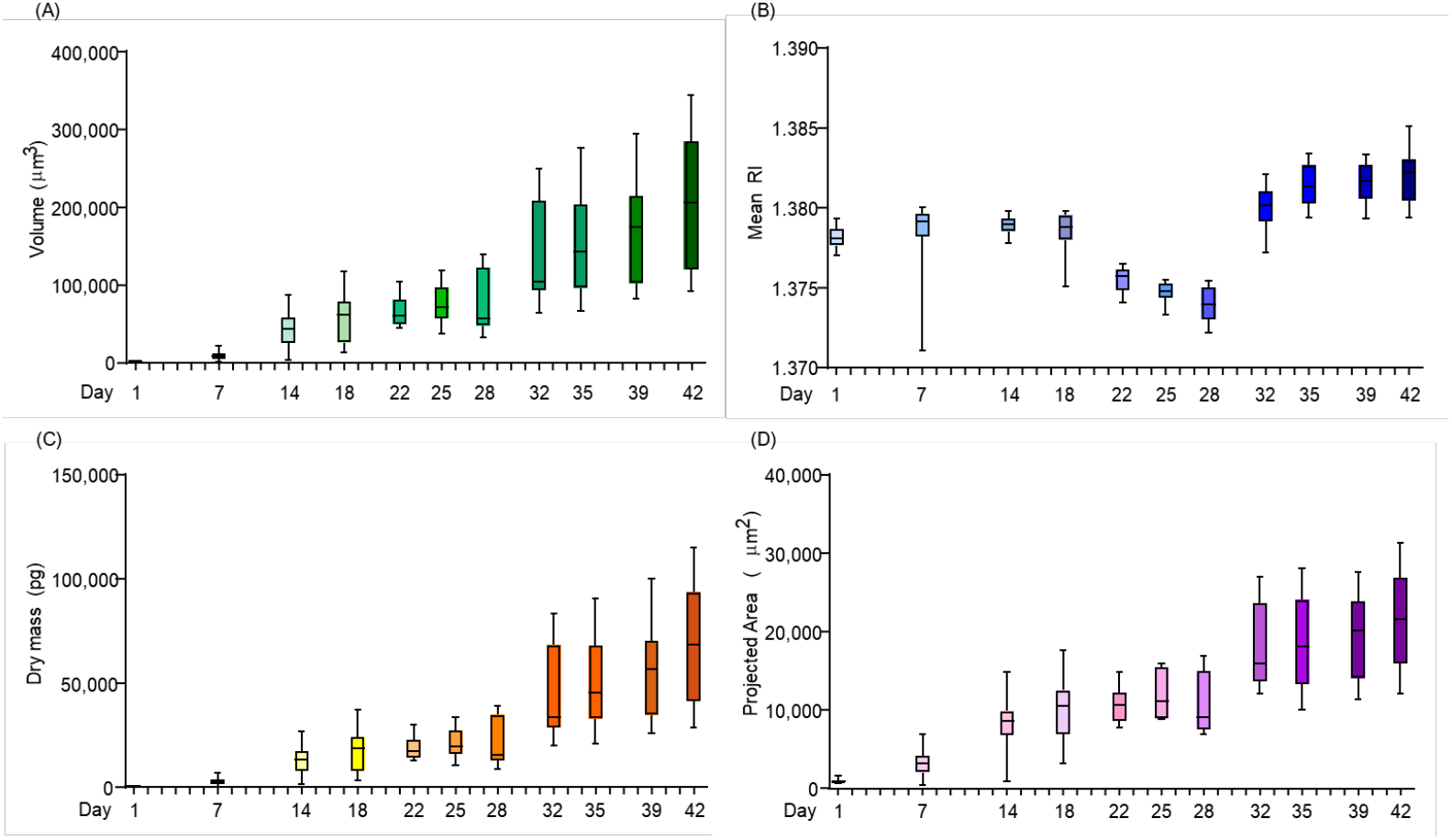
Statistical Distribution of Lipid Droplet Parameters in Adipocyte Redifferentiation Over 42 Days. (A) The volumetric expansion of lipid droplets across different time points, depicting a growth from 1,627 μm^3^ at the outset to 207,030 μm^3^ at the end of the 42-day observation period. (B) The mean refractive index of lipid droplets, indicating an incremental rise from 1.378 on Day 1 to 1.382 on Day 42, suggesting an increase in lipid content.(C) The dry mass accumulation in lipid droplets, with a marked increase from 497 pg to 65,776 pg during the 42-day redifferentiation period. (D) The increase in the projected area of lipid droplets, from 963 μm^2^ initially to 21,666 μm^2^ on Day 42, highlighting the growth in lipid droplet surface area. Each box plot represents data from 10 different measurements (N=10), providing a comprehensive view of the dynamic changes in lipid droplet characteristics during adipocyte maturation. Scale bar represents 40 μm.

Subsequently, to ensure the inclusion of smaller, developing LDs, the images underwent filtering through the Tophat method, followed by an additional thresholding step. The integration of these two procedures allowed for effective segmentation of both large and small LDs from the HT images, ensuring a thorough analysis of lipid droplet dynamics throughout the adipocyte differentiation process.

The volumetric analysis (Figure 4A) revealed a dramatic growth in the lipid droplets, expanding from an initial 1,627 μm^3^ to 207,030 μm^3^, indicative of the cells’ lipid accumulation capacity. Figure 4B showcases the mean RI increment from 1.378 to 1.382 over the 42-day period, reflecting an increase in lipid content within the droplets. Furthermore, a significant rise in dry mass from 497 pg to 65,776 pg was recorded (Figure 4C), and the projected area of lipid droplets increased from 963 μm^2^ to 21,666 μm^2^ (Figure 4D), corroborating the volumetric data. The inflection point at Day 28 and the lack of plateauing beyond Day 30 are likely due to the ongoing differentiation and maturation processes in adipocytes that extend beyond the initial phase of differentiation expected by Day 21. This prolonged activity may reflect the continued cellular and metabolic changes that occur as adipocytes mature.

These quantitative analyses confirm the visual observations from the time-lapse imaging, providing an objective measure of the lipid droplets’ physical properties. Together, the holistic view of adipocyte redifferentiation offered by Figures 3 and 4 underscores the complex nature of cellular differentiation, integrating morphological, functional, and quantitative data to construct a comprehensive narrative of adipocyte biology.

### Longitudinal Visualization of Adipocyte Redifferentiation Highlights Lipid Droplet Dynamics

To analyse adipocyte redifferentiation through longitudinal time-lapse imaging across 10 distinct microscopic fields over a period of 42 days, the morphological changes within individual adipocytes were investigated, specifically focusing on the dynamics of lipid droplet growth (Figure 5A). Quantitative tracking of lipid droplet volume revealed a consistent and significant expansion, delineating the cells’ redifferentiation trajectory towards a mature, lipid-rich phenotype (Figure 5B).

**Figure 5:**
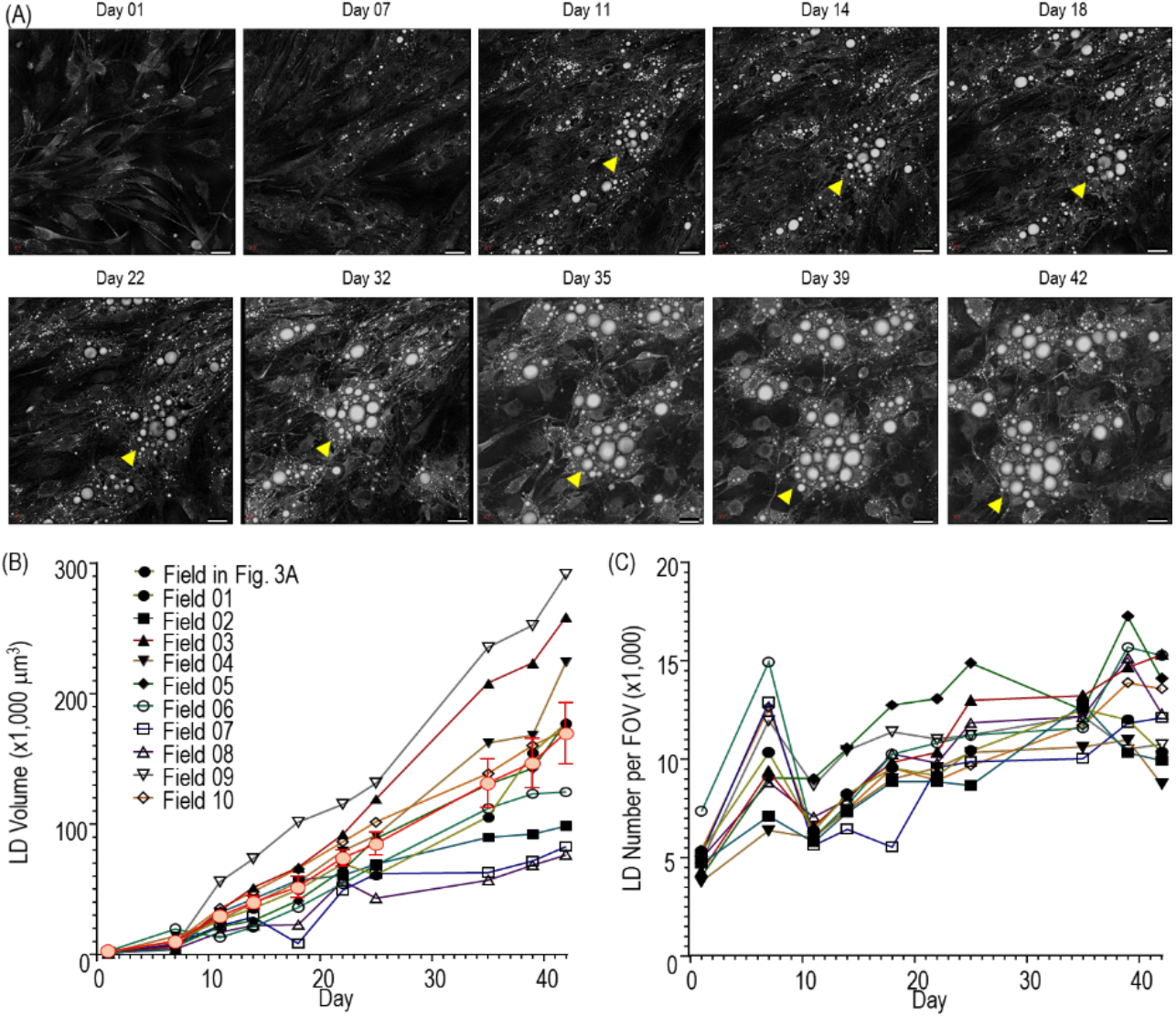
Sequential Visualization and Quantification of Adipocyte Redifferentiation. (A) Time-lapse series depicting adipocyte differentiation across 10 different microscopic fields over a 42-day period, with snapshots taken on Days 1, 7, 11, 14, 18, 22, 32, 35, 39, and 42. Yellow arrowheads indicate the same region being taken over time, pointing the gradual increase in lipid droplet size. (B) Graphical representation of the average lipid droplet volume over time, aggregated from the 10 observation fields, indicating a consistent increase over the 42-day period. Data points are expressed as mean ± SEM (the red line). Trajectory plots for each field, with each line representing the change in volume of lipid droplets for a single field, underscoring the variability and trend of growth across individual observations. (C) Trajectory plots for each field, with each line representing the number of lipid droplets for a single field. Scale bars correspond to 40 μm.

Differential growth patterns of lipid droplets across the cell population were discerned through field-specific trajectory analysis, highlighting the heterogeneity inherent to cellular differentiation processes (Figure 5C). These data not only underscore the complexity of adipocyte maturation but also validate the methodological precision of our approach.

### Longitudinal Visualization of 3T3-L1 cell Differentiation to Adiopocyte

To demonstrate the capability of measuring fast dynamics of Adipocyte differentiation during early differentiation and the applicability of the present method during the fast dynamics, adipocyte differentiation of 3T3-L1 cell was measured for 96 hours with the time interval of 1 minute (Fig. 6)

**Figure 6:**
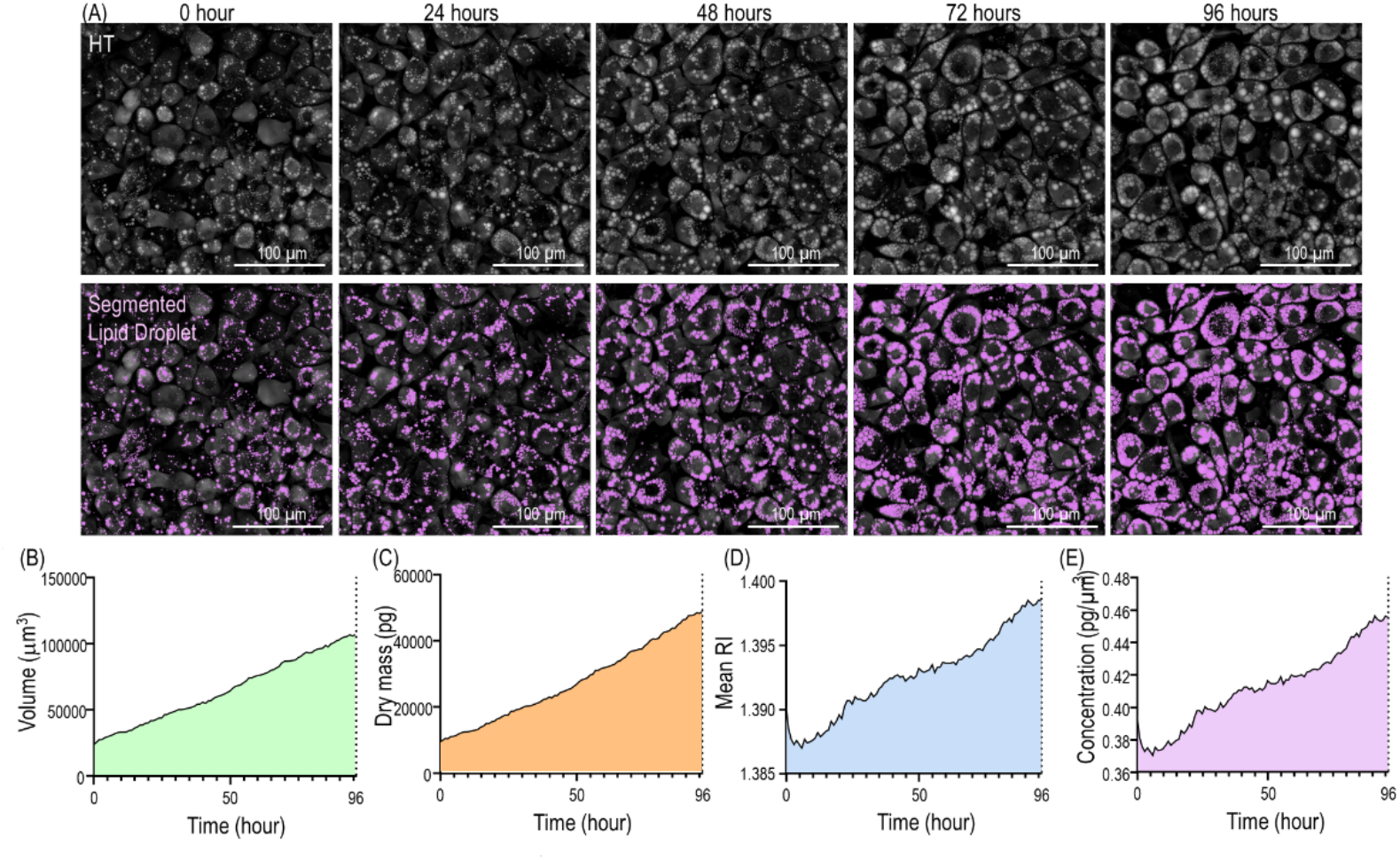
Longitudinal Visualization of Lipid Droplet Dynamics during Adipocyte differentiation. The top row shows holotomography (HT) images of 3T3-L1 to adipocytes at different stages of differentiation over a 4-day period. The middle row presents corresponding segmented lipid droplets (LDs) highlighted in purple, demonstrating the changes in lipid droplet morphology and distribution as adipocytes mature. The bottom row includes quantitative measurements representing the growth trends in LD volume (green), dry mass (orange), mean refractive index (blue), concentration (purple) over time. Scale bars in each image represent 100 μm. This figure illustrates the progression of adipocyte differentiation and the associated biophysical changes in lipid droplets.

The result presents a comprehensive visualization of the LD dynamics during the differentiation process of adipocytes over a 4-day period. Figure 6(A) the sequential stages of adipocyte redifferentiation captured using HT, revealing the cellular morphology with high resolution. These images illustrate the progressive changes in the structure of the adipocytes as they transition from their initial state to more mature stages, with notable increases in the size and density of the lipid droplets. Figures 6(B)-6(E) effectively demonstrate how holotomography can be used to monitor the redifferentiation process in adipocytes, providing both qualitative and quantitative insights into lipid droplet dynamics. The combination of high-resolution imaging and precise segmentation allows for a detailed understanding of the progression of adipocyte maturation, which is critical for exploring the mechanisms of lipid storage and mobilization in metabolic research.

## 4. DISCUSSION

The method we introduce offers an approach to the label-free long-term observation and real-time tracking of the redifferentiation process in DFAT cells, utilizing HT and advanced quantitative image analysis. This approach provides unique insights into the intricate subcellular dynamics over an extended redifferentiation timeline, a capability not readily achievable with existing methodologies.

Recent advancements in morphological analysis of LDs have facilitated the quantification of both macroscopic and microscopic changes within adipocytes, linking these changes to lipid production [27, 28]. However, these methods often fall short in their ability to monitor live cells over extended periods. The use of fluorescent dyes, while enabling live cell observation, poses challenges for long-term tracking due to dye persistence and potential phototoxicity. Our technique effectively overcomes these limitations, bypassing the need for traditional staining methods that can interfere with cellular functions and hinder continuous observation. This ability to conduct uninterrupted, long-term tracking without the interference typically associated with staining protocols marks a significant advancement in the field of cellular biology and adipocyte research.

The novelty of our study lies not only in using HT technology but also in applying this technology to investigate the long-term differentiation process of adipocytes derived from patient-specific fibroblasts, particularly in the context of KOA. This approach provides new insights into lipid dynamics within human disease contexts, which has not been extensively explored using this technology.

The refined integration of morphological assessments with quantitative data has furnished a comprehensive framework for exploring adipocyte biology. A marked correlation between the morphometrics of LDs and the expression profiles of adipogenic genes elucidates the complex processes of adipocyte redifferentiation. These findings enrich our understanding of lipid storage mechanisms within adipocytes, shedding light on the broader principles of cellular differentiation amid metabolic activity and disease progression.

While the HT instrument itself is commercially available, the innovative aspect of our study lies in the custom quantitative imaging algorithms we developed. These algorithms enable precise segmentation, tracking, and quantitative analysis of lipid droplets over extended periods, which is crucial for understanding adipocyte maturation and its role in inflammation and metabolic changes associated with KOA.

Furthermore, our approach integrates high-resolution 3D imaging with biological assays, such as immunofluorescence and gene expression analyses, to provide a comprehensive view of adipocyte behavior. This integrative method is novel in its ability to correlate changes in lipid droplet morphology and distribution with genetic markers of adipocyte differentiation and inflammatory responses in KOA.

In particular, the enlargement of LDs as captured using HT corresponded with a significant upsurge in the expression of adipogenic markers such as LPL. This correlation emphasizes the reliability of LD morphology as an indicator of adipocyte maturation. The pronounced correlation between the observed cellular alterations and molecular indicators of adipocyte function highlights HT’s capability in delineating the subtle phases of cell maturation. This, in turn, provides novel insights into the intricate relationship between cellular structure and function throughout the maturation of adipocytes.

Our study’s quantitative analysis extends beyond mere observation, providing insight into the biophysical changes that occur during adipocyte redifferentiation. The calculated volumes, mean RI, dry mass, and projected areas of individual lipid droplets form a detailed profile of adipogenesis at the subcellular level. These metrics not only facilitate an understanding of lipid storage and mobilization within the cells but also serve as potential biomarkers for monitoring and evaluating the efficacy of therapeutic interventions aimed at modulating adipocyte differentiation and function.

The heterogeneity observed in the growth patterns of lipid droplets underscores the individual variability in cell behavior during the redifferentiation process. This variability could have significant implications for the pathogenesis of KOA, as it suggests that individual adipocyte responses within the IFP may contribute differently to the disease state. Our findings, which enable the tracking of dynamic changes in lipid droplets and gene expression, could pave the way for targeted therapeutic strategies that address adipocyte function within the joint. The study also opens avenues for personalized medicine approaches in KOA treatment, tailored to the unique cellular responses of the patient’s adipocytes.

We have emphasized the utility of applying HT technology to study the role of adipocytes in disease contexts, particularly in inflammatory conditions like KOA. This approach has potential implications for developing targeted therapies that address metabolic and inflammatory pathways in adipose tissues within the joint environment.

Looking forward, the potential for HT imaging to be applied to other adipocyte related pathophysiological studies is immense. The present approach also holds promise for investigating lipogenesis and lipid metabolism processes across various cell types [42, 43, 60]. Additionally, integrating machine learning techniques with HT could further enhance the analysis [61-63], potentially automating the recognition and quantification of cellular changes and classification of cell types based on LD contents. The automatic detection and virtual staining of major subcellular organelles as well as lipid droplets would expedite the systematic long-term label-free assessments of various metabolisms and lipodomics [64-66]. Integrating HT with other imaging techniques, such as Raman spectroscopy and fluorescence imaging, could enhance our understanding by offering additional insights into the types of lipids present and their molecular specificities [60, 67, 68]. This correlative approach promises a more comprehensive analysis, combining the structural and quantitative strengths of HT with the molecular-level detail provided by Raman and fluorescence imaging. Additionally, the current methodology can be integrated with a microfluidic platform, thereby facilitating high-throughput analysis of unlabeled live cells [69, 70]. This combination would significantly enhance the efficiency and scale of cellular analysis, broadening the potential applications in live cell studies. Such advancements could revolutionize applications in cancer biology [71], 3D biology and organoids [72, 73], tissue engineering [74], regenerative medicine [75], and drug discovery [76], extending well into the study and treatment of various metabolic diseases beyond KOA.

In conclusion, this study has provided substantial advancements in our understanding of adipocyte biology, particularly in the context of cell redifferentiation. The insights garnered from our holistic approach underscore the intricate balance between cellular structure and function during adipocyte maturation. The implications of our work extend to the pathology and treatment of KOA, with the promise of broader applications that could significantly impact the fields of regenerative medicine and metabolic disease treatment.

## Funding

National Research Foundation of Korea (2015R1A3A2066550, 2022M3H4A1A02074314); Institute of Information & communications Technology Planning & Evaluation (IITP; 2021-0-00745) grant funded by the Korea government (MSIT); KAIST Institute of Technology Value Creation; Industry Liaison Center (G-CORE Project) grant funded by MSIT (N11230131); the Korea Health Technology R&D Project through the Korea Health Industry Development Institute (KHIDI), funded by the Ministry of Health & Welfare, Korea (HI21C0977).

## Disclosures

Hye-Jin Kim, Sumin Lee, and YongKeun Park have financial interests in Tomocube Inc., a company that commercializes holotomography instruments. The other authors declare no competing interests.

## Data availability

Data underlying the results presented in this paper are not publicly available at this time but may be obtained from the authors upon reasonable request.

## Notes

### Summary of Updates

The introduction has been enhanced to include the motivation behind the study and highlight its novelty. Figure 2 has been refined by incorporating comparisons of the segmentation results with fluorescence imaging. The subsection titled "Longitudinal Visualization of 3T3-L1 Cell Differentiation to Adipocyte" and Figure 6 have been added. The discussion section has been expanded and improved.

